# A database of over 15.000 strain design publications reveals a conserved set of metabolic engineering targets across microbial hosts and products

**DOI:** 10.64898/2025.12.15.694291

**Authors:** Elisa Márquez-Zavala, Francesca Di Bartolomeu, Daniel Machado

## Abstract

Microbial biotechnology has the potential to address several societal issues through the sustainable production of industrially relevant compounds. Despite decades of successful cases, rational engineering of microbial metabolism is still a complex process due to the fine balance between nutrient supply, allocation of cellular resources, energy demand and redox balancing. In this work, we implemented a text-mining workflow for metabolic engineering and compiled a database of experimentally validated strain design strategies from over 15.000 research articles, which includes information on host strain, target compounds, and gene modifications. This large dataset reveals trends on the selection of suitable hosts for different kinds of products and the respective gene targets. Despite the wide variety of microbes and products, we observe a conserved set of target metabolic genes associated with central carbon metabolism, especially in upper glycolysis, pentose-phosphate pathway, citric acid cycle, and fermentative pathways. The most distinguishing feature among strain design strategies seems not to be which genes are targeted, but rather the direction in which they are modified (increased or decreased expression). Controlling flux at key branching points and redox balancing reactions is thus a critical engineering step to steer metabolism. Our collection of 25 years of literature can provide a stepping stone for starting new strain design projects without reinventing the wheel.

## Introduction

Metabolic engineering uses genetic and molecular techniques to manipulate cellular metabolism and achieve specific goals such as the synthesis or breakdown of target chemical compounds ^1,2^. This manipulation has played an important role in sustainable bioprocessing, including the production of biofuels, bioplastics, and pharmaceuticals ^1–4^, offering solutions for challenges in medicine, agriculture, and environmental conservation ^5–8^. Despite its significant contributions, metabolic engineering encounters challenges that require innovative solutions. Among these challenges are: maintaining a balance between product yield and cell viability, managing complex cellular responses to genetic changes, and effectively scaling up bioprocesses ^3,9,10^. Continuous progress in testing, designing, and optimizing metabolic pathways and strains is essential to tackle these challenges ^9,11–13^.

The process of metabolic engineering begins with extensive knowledge of metabolic and regulatory pathways specific for the selected organisms and desired compounds ^9,11,12,14^. A combination of expert knowledge, *a priori* insight, and (*in silico*) rational strain design methods can be applied to find target genes for expression regulation, deletion, or heterologous insertion ^1,10,11,15–17^. Once gene targets are identified, synthetic biology tools (such as the CRISPR/Cas9 system) are employed to implement such designs *in vivo* ^1,11,18^. Naturally, one would select a host organism that already performs the desired task (synthesis or degradation of a target compound). Databases of functionally annotated genomes can be searched for the presence of relevant enzymes ^4,19–22^. However, the respective organisms might not be easily cultivated in simple media, might suffer from slow growth rates, or the synthetic biology toolbox might not yet be available. The selection of host organisms therefore often falls on the popular *workhorses* of industrial biotechnology, such as *Escherichia coli* and *Saccharomyces cerevisiae*.

The typical workflow of a metabolic engineering project comprises multiple interdependent decisions, such as the selection of substrates and products (based on techno-economic and life-cycle analysis) and the selection of suitable organisms and necessary genetic manipulations (based on genomic and physiological constraints). These decisions are often based on biological insight gathered from previous projects, the scientific literature, and data scattered across different databases. One strategy to address these challenges is to create a comprehensive database containing information about enzymes, pathways, organisms, products, and their relationships ^4,12,14,23^. Such a resource could simplify the process of designing and discovering new bio-based products more efficiently ^4,24,25^. One obvious approach is to leverage all the information available in the scientific literature. Databases like NCBI PubMed and Europe PMC ^26,27^ provide data from scientific articles and books, and can be accessed programmatically ^28–31^. Although these databases do not directly compile specific information such as products, genetic modifications, or organisms, additional resources can be used to develop tools for extracting and analyzing such data. Databases like ChEBI ^32,33^, UniProt ^20^, and KEGG ^19,34,35^ contain detailed information about genes, organisms, and biochemical products that can be used to further annotate and contextualize these data.

LASER (Learning Assisted Strain EngineeRing) ^24,25^ was the first specialized database focusing on strain designs extracted from the literature. It was manually curated and contains entries from hundreds of articles for two well-known organisms (*E. coli* and *S. cerevisiae*). This manual curation process, while ensuring accuracy, also limited the size, scope, and maintainability of the database. This is where automatic extraction tools can be highly useful ^36^. In this work, we used LASER to train a machine learning classifier and created an updated strain design database using automatic extraction from over 15.000 articles available on PubMed. The scope is expanded to cover a larger number of organisms and to account for a decade of new scientific literature. We show how this database can be used to extract insight on associations between host organisms, target products, and the frequently targeted metabolic pathways.

## Methods

### Extracting information from the LASER database

Data from the LASER database ^24,25^ was retrieved by accessing all the files on the respective repository and parsing them using Python scripts. We extracted relevant information such as organisms, products, genes, and pathways.

### Finding relevant articles from the literature

Relevant articles were identified by programmatically searching the PubMed database using the NCBI E-utilities API ^30^ accessed via the Biopython Entrez module (Biopython v1.85) ^37^. To construct the search queries, we assembled a list of keywords related to metabolic engineering and production improvement. These included stemmed forms with an asterisk (e.g., improv* or engineer*), which allows PubMed to match multiple word variants (e.g., improve, improved, engineering, engineered).

The search terms were combined into structured queries and applied to articles published between 2000 and 2025. PubMed IDs (PMIDs) returned by these searches were collected, and for each article the corresponding metadata (title, abstract, journal, publication year, authors, DOI, and PubMed Central ID) was retrieved through the API. We also compiled a list of journals to exclude based on quality and relevance to the topic. Articles were preprocessed by stop-word removal, lowercasing, stemming, and normalization of organism and compound names. To mitigate the effect of highly frequent or generic terms, we applied TF–IDF ^38,39^ weighting, which down-weights common words and reduces representation bias.

The articles were then classified as relevant or not using a supervised approach. For this, we constructed a labeled dataset consisting of a positive and a negative set. The positive set included articles from journals dedicated to metabolic engineering (e.g., Metabolic Engineering, Microbial Cell Factories) as well as all curated LASER database articles. For the negative set, we generated a list of journals exclusive to other research fields, based on domain-specific exclusion keywords. A manually curated set of exclusion terms (e.g., oncology, botany, neuro, clinic) were matched against the National Library of Medicine (NLM) journal catalog ^29^, and all journals whose titles contained any exclusion term were collected into an exclusion list. We then executed the same PubMed search strategy used for retrieving potentially relevant articles but restricted it to this exclusion list of journals. The resulting articles, originating from journals outside fields likely to contain metabolic engineering work, formed the pool for the negative set. From this pool, we randomly sampled articles without replacement to obtain a negative set matching the positive set in size.

We evaluated several text-classification models, including Bernoulli Naive Bayes, Multinomial Naive Bayes, Random Forest, and Support Vector Classifiers. For each model, we trained three input configurations: title only, abstract only, and a combined title and abstract. Model performance was assessed using LASER articles. Based on this evaluation, four models were selected as base learners for a stacked ensemble: two linear Support Vector Classifiers trained on the combined input, a Multinomial Naive Bayes model trained on the combined input, and a Multinomial Naive Bayes model trained on abstracts only. Their output probabilities were combined using a logistic-regression meta-classifier, which served as the final system for all downstream classification tasks.

### Extracting full text

After classification and final selection of relevant articles, full texts were retrieved using two complimentary approaches. For articles indexed in PubMed Central, XML full texts were downloaded using the NCBI E-utilities API ^30^ via the Biopython Entrez module (Biopython v1.85) ^37^. For articles without PMC access, open-access PDFs were located through the Unpaywall REST API (Unpaywall v0.2.3) ^40^ and downloaded accordingly. PDFs were converted to text using PDFMiner.six v20250506 ^41^. All extracted texts underwent standardized post-processing to remove metadata, normalize formatting, and retain only the main article body.

### Extracting organism names

Organism names were extracted from the main text of the articles using the NCBI taxonomy database. We filtered out non-relevant organism names, such as viruses, environmental samples, mixed libraries, or miscellaneous sequences. We normalized the text of the articles by applying lowercasing, punctuation removal, and whitespace trimming, and searched for the name of the organisms in the text using regular expressions and string matching. For each match, we extracted the scientific name and taxonomic rank of the organism.

### Extracting product names

Products were extracted from the articles by applying natural language processing. We simplified and shortened the sentences by splitting them into clauses and removing adjectives and adverbs that did not affect the meaning of the sentence. We performed part-of-speech tagging and dependency parsing on each sentence using spaCy (3.7.2) ^42^. We identified potential products as nouns or noun phrases that were modified by verbs such as “produce”, “synthesize”, or “enhance”. We standardized the product names using the ChEBI database ^32,33^, and also standardized the notation of stereochemistry and chirality (by replacing *R, S, L, D* with their respective symbols).

### Extracting gene names and orthology

Gene names were extracted from the text using organism-specific information. First, we downloaded all gene names mentioned in UniProt ^20^ for the organism’s name(s) found in each article. These were then searched in the article text by string matching. To standardize the gene names, and facilitate comparison across organisms, we mapped them to KEGG Orthologs (KO), either directly from the respective cross-reference UniProt (if available), or indirectly, through a query on the KEGG database ^19^.

### Mapping to Enzyme Commission (EC) numbers and BiGG reactions

In addition to KO mapping, we also retrieved Enzyme Commission (EC) numbers from UniProt cross-references ^20^. These EC numbers were then used to link reactions across different organisms, by querying the MetaNetX database ^43^, which integrates data from multiple sources, including BiGG Models ^44^. MetaNetX allowed us to identify the corresponding BiGG reaction identifiers for each EC number. This enabled pathway-level visualizations where frequent genetic interventions across organisms could be compared within the same metabolic context.

### Extracting gene modifications

Gene modification events were identified using natural language processing. The text was segmented into sentences and clauses, and part-of-speech tagging and dependency parsing were applied with spaCy (3.7.2) ^42^ to detect actions related to genetic changes. We searched for verbs commonly associated with genetic engineering (e.g., delete, disrupt, mutate, knock out, overexpress) and linked them to previously identified gene names. Noun-phrase patterns such as “deletion of gene” or “mutation of gene” were also captured. All detected terms were normalized to a controlled vocabulary for consistency across articles

Normalized modifications were then classified into *Positive, Negative*, or *Other* using a curated rule-based lexicon. Terms indicating increased activity or expression (e.g., overexpress, upregulate, enhance) were labeled *Positive*, while those implying reduced function or loss of activity (e.g., delete, knock out, repress, inhibit) were labeled *Negative*. Ambiguous terms (e.g., mutate, engineer, regulate) were assigned to *Other*. Negation patterns (e.g., “not overexpress”) were detected to reverse polarity, with negative cues taking precedence when conflicts occurred.

## Data availability

All the source code and data generated in this work is publicly available in the following repository: https://github.com/emarquezz/ELISER-StrainDesignDB.

## Results

### Literature mining yields a collection of over 15,000 relevant articles

The initial literature search yielded approximately 150,000 articles containing terms related to metabolic engineering. Upon inspection, we observed that this initial set included several false positive results, including articles and journals not relevant to the topic. Therefore, we compiled a list of journals to exclude from further analysis and we trained different classification algorithms to further filter our collection. We created a training dataset using articles in the LASER database as a positive set and articles from unrelated journals as a negative set. We evaluated the impact of training the classifiers using only the article titles, abstracts, or both. The hyperparameters used in each classifier are available in the supplementary material.

Looking at the trade-off between specificity and sensitivity for the different classifiers (Supp. Fig. 1), we observe that, despite a similar performance, the Multinomial Naive Bayes (MNB) classifier displayed the best overall performance, which is consistent with previous literature citing this method as effective for text classification ^45–47^. We can also observe that using the abstract or both abstract and title, results in a better performance than training on title alone. Therefore, we used the classification results of MNB using both title and abstract as the final dataset. This reduced the initial list of 152,921 articles to 20,108. Through additional inspection, we observed that about 20% of these were in fact review articles, which we excluded from further analysis, reducing the dataset to 16,130 articles.

**Figure 1.**
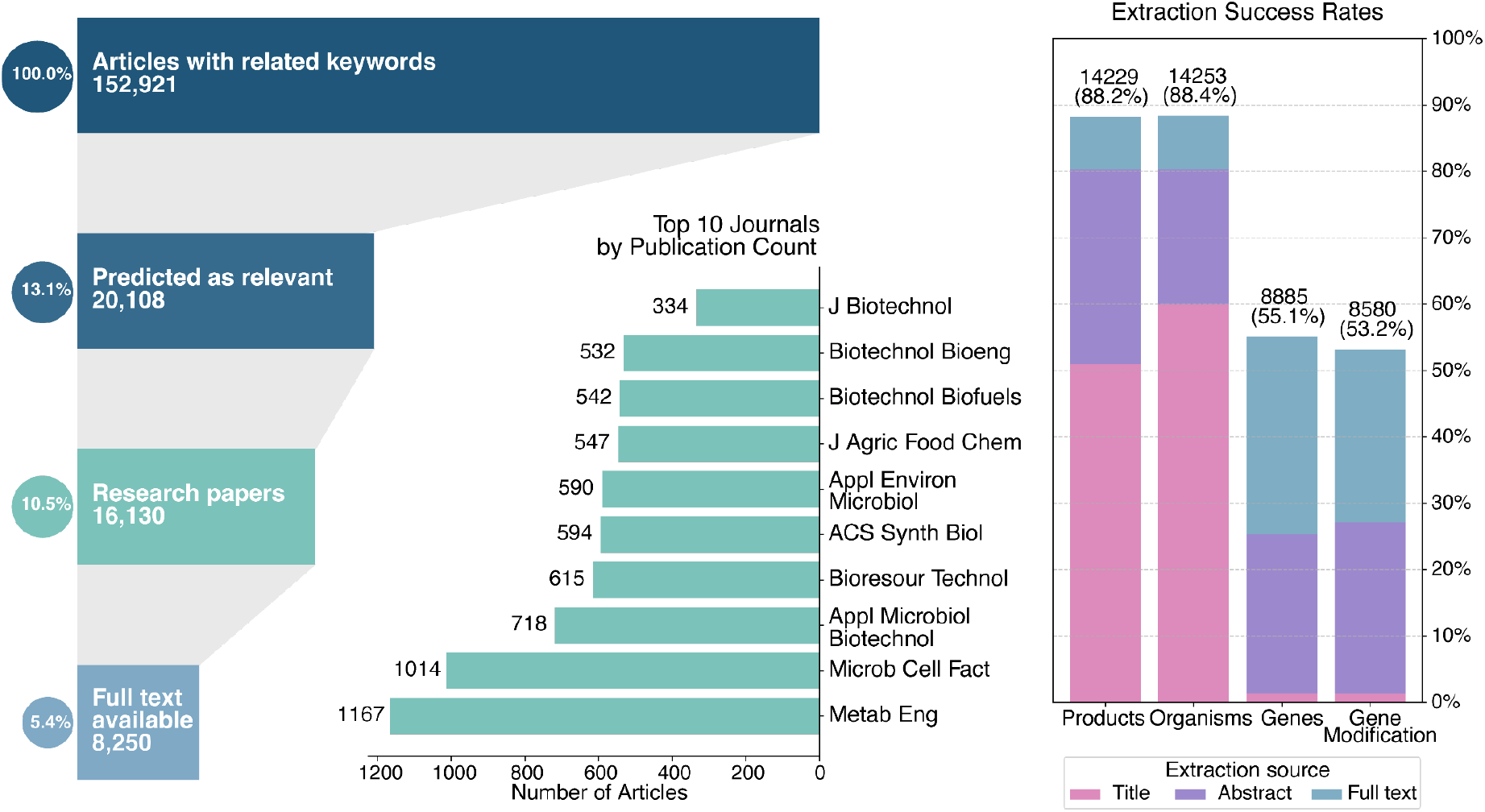
Total information obtained at each step in our workflow. (Left) Funnel plot showing progressive filtering: initial PubMed search, machine learning classification, removal of review articles, and full-text availability. (Middle) Top 10 journals in the final dataset. (Right) Extraction success rates for products, organisms, genes, and gene modifications by source (title, abstract, full text).

We then proceeded to extract relevant information from the articles, including organisms, products, and target genes. Since many articles did not describe the modified genes either in the title or abstract, we tried to obtain the full text of all the articles classified as relevant (see Methods) and were able to download approximately 50%. For the remaining articles, we proceeded with extracting information only from the title and abstract. After accounting for articles where extraction was successful, the final dataset comprised 15,627 articles. Overall, we were able to extract the organism and product for most of the articles, and the modified genes for about half of the articles. Figure 1 shows an overview of the amount of information retained at each step in our workflow.

### Trends in the utilization of industrial organisms

We used the NCBI Taxonomy database to extract and normalize organism names. We first tried to identify the main organism mentioned in each article, but additionally retained all mentioned organisms, which facilitated, at a later stage, the identification of genes from heterologous pathways. To streamline the search process, we prioritized titles, followed by abstracts, and finally the full text. After normalizing organism names at species level, we identified 1,352 species across 702 genera. As expected, the species frequency shows a very skewed distribution (Fig. 2a), with the two main industrial workhorses, *E. coli* and *S. cerevisiae*, comprising more than 40% of all the entries.

**Figure 2.**
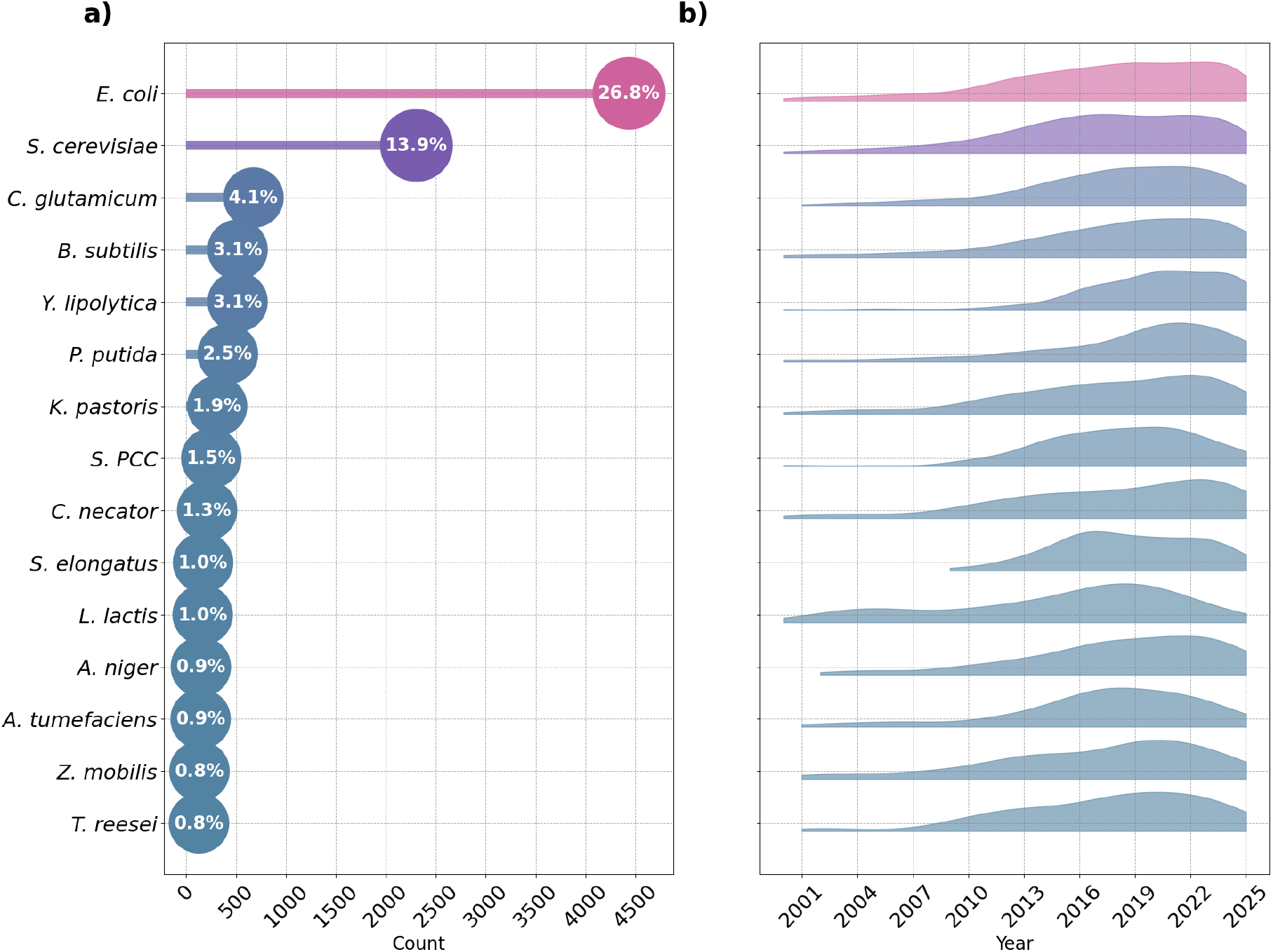
Summary of the most frequent (top 15) organisms: a) frequency distribution of each species in the final dataset; b) temporal evolution of the number of citations for each species.

We also analyzed trends in the utilization of each organism in a 25-year period (2000-2025). We can see that, while *E. coli and S. cerevisiae* have been constantly used throughout the years, other organisms, such as *Yarrowia lipolytica* and *Pseudomonas putida*, show a recent increase in citations. This indicates an emerging interest in using a wider variety of organisms as microbial cell factories.

### A catalog of microbially produced molecules

Identifying the target products in the text revealed to be more challenging than finding organisms. We used ChEBI to obtain compound names and synonyms. However, to distinguish between the target product and other compounds (substrates or other intermediates) mentioned in the text, we employed Natural Language Processing (NLP) techniques to semantically analyze sentences such as “production of *x*” (see methods). After extraction, we standardized synonyms and alternative spellings using the ChEBI ontology, which resulted in approximately 4,128 unique products. This classification, however, varies in granularity, with some products corresponding to specific compounds (e.g.: lactate, succinate) and others corresponding to compound classes (e.g.: protein, lipid).

Looking at the distribution frequency of products, we can observe that it is not as skewed as the organism distribution (Fig. 3a). Biofuels (when grouped together) are the common target products (12.7% of all references), followed by amino acids, carotenoids, and proteins.

**Figure 3.**
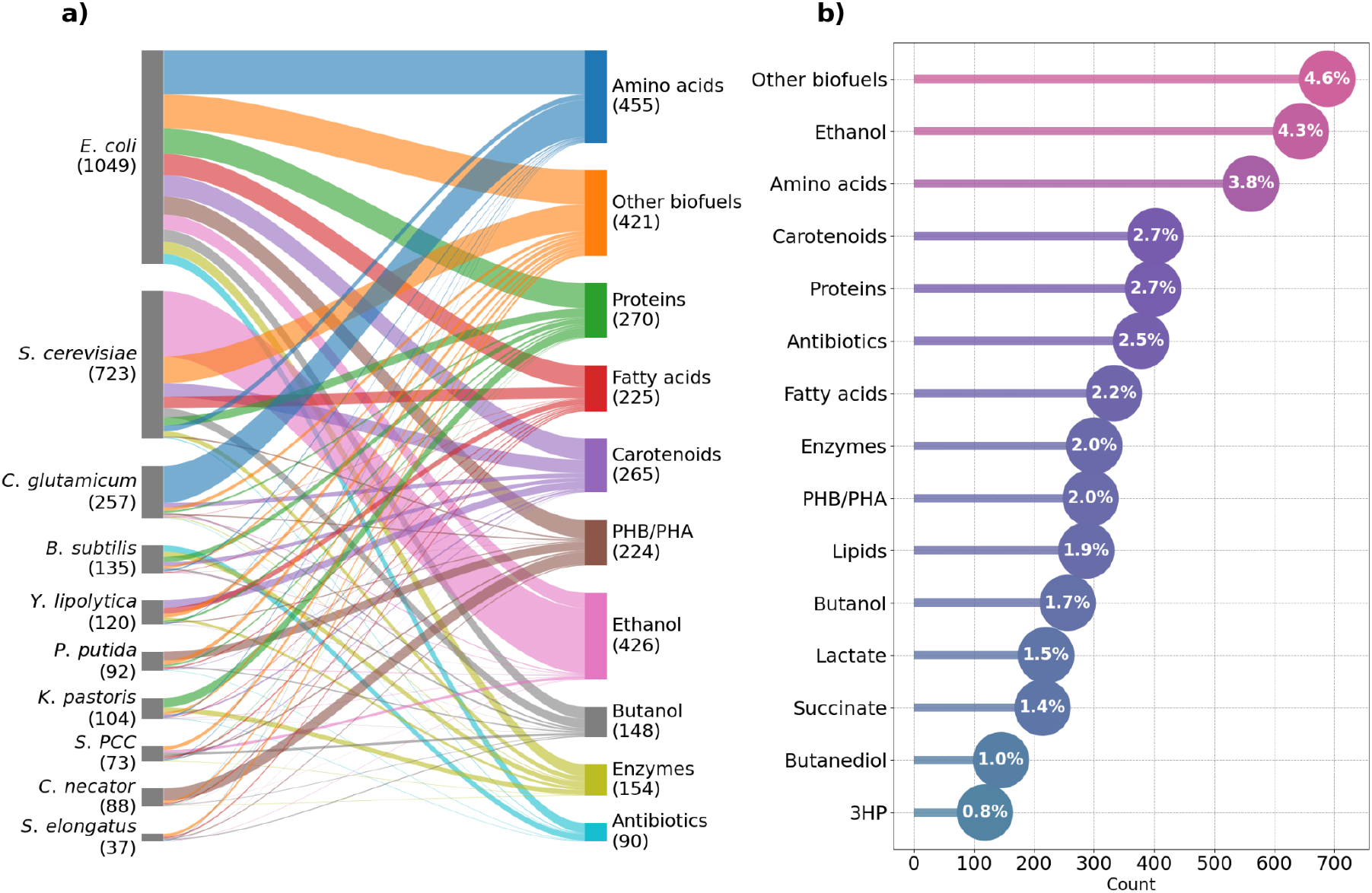
Analysis of most common target products: a) frequency distribution of the top 15 products; b) Sankey diagram representing the association between the 10 most common organisms and 10 most common products.

To understand preferences between selected hosts and respective target compounds we analyzed their co-occurrence (Fig. 3b). We can observe that classical workhorses like *E. coli* and *S. cerevisiae* have a broad range of utilization and have been used to produce all kinds of compounds. On the other hand, it is possible to observe the utilization of *specialist* hosts like *Komagataella pastoris* (previously known as *Pichia pastoris*), mainly used in protein production, *Cupriavidus nector* mainly used for the production of polyesters, and *Corynebacterium glutamicum*, standing out as the preferred host for the production of individual amino acids.

### A common subset of genes is consistently targeted

The extraction of target genes posed additional challenges due to many gene names (e.g: *gap*) being also valid English words. Therefore, we used UniProt to extract all possible gene names and synonyms, limited to the organisms mentioned in each article (see methods). To standardize gene identifiers and to facilitate comparison of frequently targeted genes across organisms, we mapped all organism-specific genes into KEGG orthologs. In addition, we also linked these genes to EC numbers via UniProt and further mapped them to BiGG reaction identifiers. This allowed us to link genetic interventions to corresponding reactions and pathways, providing a unified metabolic context across organisms. We successfully extracted gene-related information from 8,885 articles. In addition, we used NLP to find verbs associated with these genes (such as “express” or “regulate”) and grouped the genetic interventions into three categories: *positive* (including up-regulation, increase in copy number, or heterologous insertion), *negative* (including knockout, down-regulation, or deleterious mutations) or *other* (including unspecific actions such as: target, modify, regulate).

We can observe that the vast majority of frequently targeted genes are associated with reactions in central carbon metabolism (Fig. 4). Not surprisingly, the most frequently targeted area revolves around the pyruvate node, a key branching point of metabolism, distributing flux towards respiratory and fermentative pathways. Flux input into the TCA cycle, the main factory of most cellular precursors and source of reductive potential for oxidative phosphorylation, seems to be mainly targeted through phosphoenolpyruvate carboxylase (PPC) and citrate synthase (CS). The flux directed towards fermentative pathways, responsible for redox balance through NAD+ regeneration, can be channeled into the desired fermentation product (lactate, acetate, ethanol) by selectively targeting any of the reactions in these pathways. Another commonly targeted area is the branching point between upper glycolysis and the pentose-phosphate pathway, which can control the production of NADPH, an essential cofactor for biosynthetic pathways, and an entry point to produce aromatic amino acids, which are precursors of many commercially relevant compounds.

**Figure 4.**
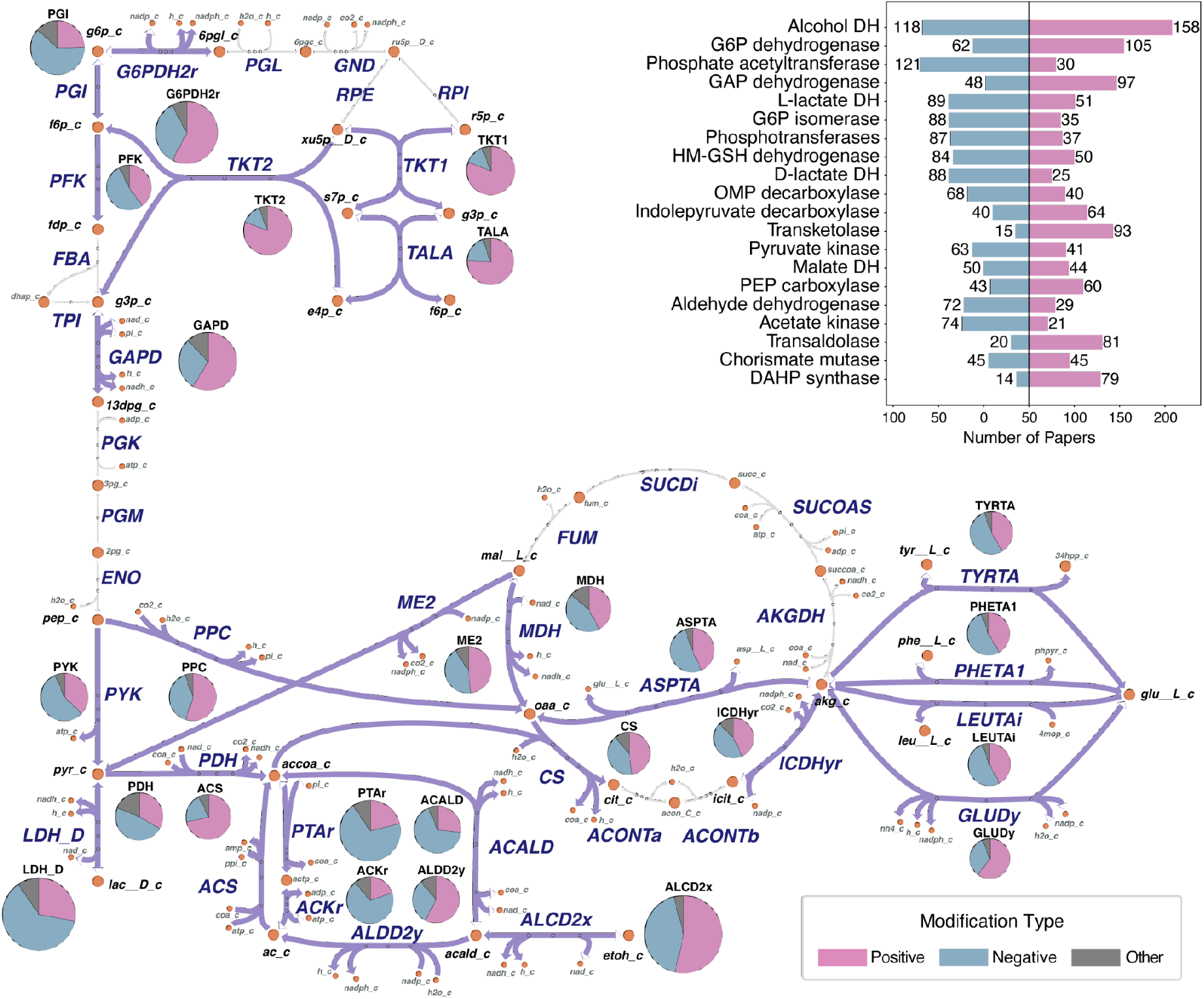
Distribution of the most frequently targeted genes and their respective reactions. The pie charts represent how often a reaction is targeted (circle size) and the type of modification performed (pink: over-expression, blue: under-expression, grey: unclassified). The bar chart (top right) shows the most frequently mentioned enzyme types, with bars colored according to the same scheme. Figure produced with a customized version of Escher-Trace ^48^.

Looking at the type of interventions performed, we can observe that there isn’t a single gene or reaction that is consistently up or down-regulated. In the most extreme cases, some reactions are targeted for up-regulation (transaldolase and transketolase) or down-regulation (phosphate acetyltransferase) about 80% of the time. Some reactions at key branching points, like the first step in the pentose phosphate pathway (glucose-6-phosphate dehydrogenase) or the TCA cycle (citrate synthase), are up or down regulated with nearly the same frequency. Overall, while there appears to be a conservation of targeted genes across applications regardless of the microbial host or target product, the direction in which the genes are manipulated is one of the main distinguishing factors among strain design strategies.

### Co-occurring modifications reveal common design strategies

Finally, we analysed the co-occurrence of genes across the literature to find groups of genes that are frequently modified together. After calculating the co-occurrence between all gene pairs in our dataset, we filtered for genes that co-occur in at least 10 articles with a Jaccard index of at least 0.2. This resulted in 9 gene clusters exclusive to either *E. coli* or *S. cerevisiae* (Fig. 5).

**Figure 5.**
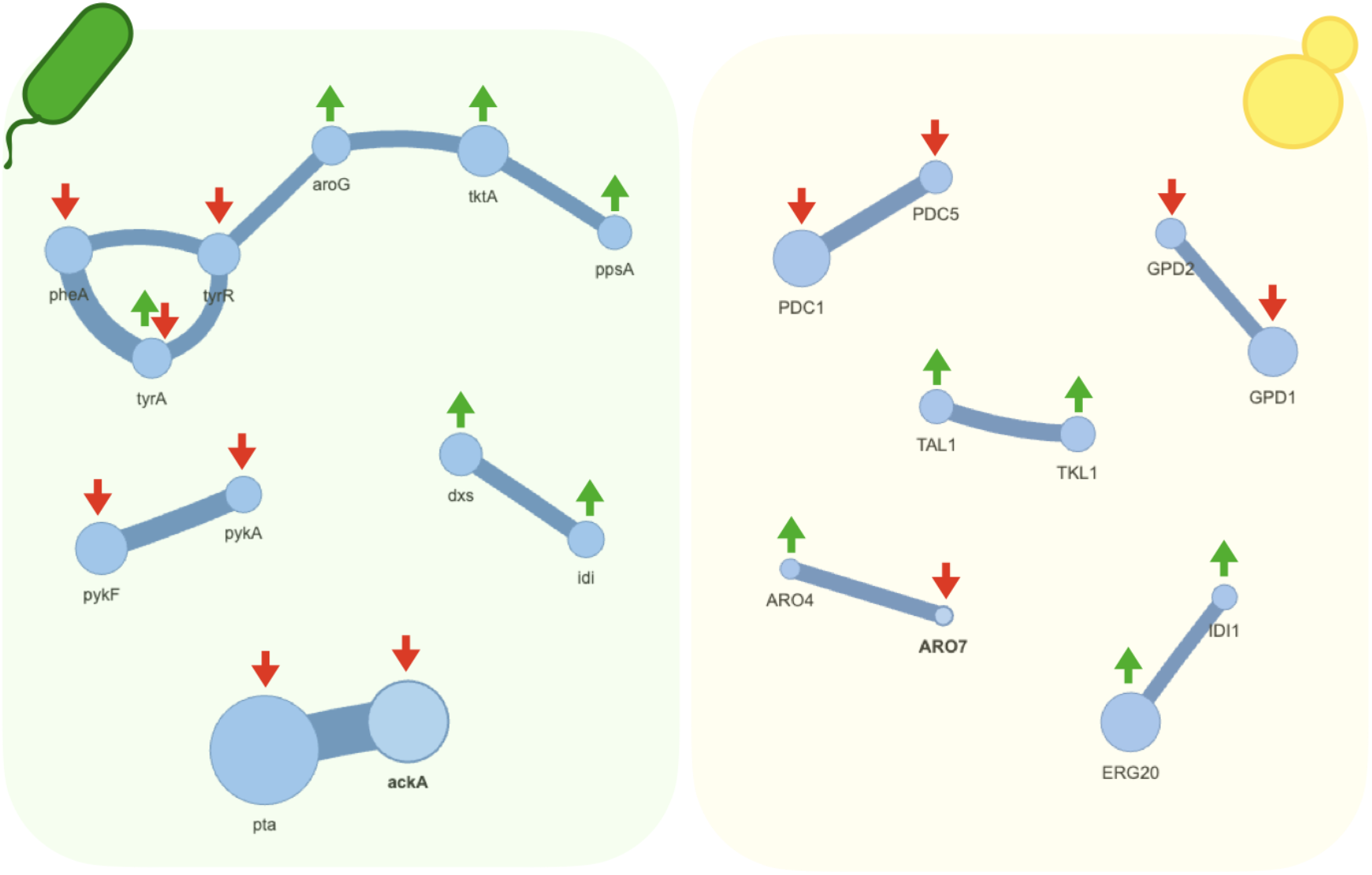
Gene co-occurrence network. Node size indicates how frequently each gene occurs in the literature and edge width indicates how frequently two genes co-occur. The up and down arrows indicate the type of modification most commonly used in the strain designs where co-occurrence was detected. The most frequent co-occurring groups are specific to either *E. coli* (left panel) or *S. cerevisiae* (right panel).

All clusters contain only two genes, except for a large cluster with 6 genes related to the production of aromatic amino acids (AAAs) in *E. coli*. The AAA pathway, also referred to as the *shikimate pathway*, is the precursor to a wide variety of industrially relevant compounds. This gene cluster includes the up-regulation of *ppsA, tktA*, and *aroG*, which increases the availability of phosphoenolpyruvate (pep) and erythrose-4-phosphate (e4p) the two substrates of the first committing step of the AAA pathway. It also includes the down-regulation of *tyrR*, the main regulatory repressor of this pathway, and the up/down-regulation of tyrA and pheA, the final branching points that split the metabolic flux into the three aromatic amino acids (phenylalanine, tyrosine, and tryptophan).

Some of the gene pairs correspond to the knockout of isozymes that catalyze the same reaction (e.g.: pykA/pykF, PDC1/PDC5, GDP1/GDP2). However, most gene pairs encode enzymes catalyzing different, but interconnected, reactions. For instance, the knockout of *pta* and *ackA* in *E. coli* blocks two consecutive reactions catalyzing the conversion of acetyl-CoA into acetate. This strategy has been used to redirect the loss of carbon through acetate secretion into other products, mainly biofuels and succinate. The up-regulation of *dxs* and *idi*, which catalyze the first and final steps of the non-mevalonate pathway, has been applied in the production of a class of secondary metabolites known as terpenes (especially carotenoids and diterpenes).

Most of the co-occurring genes found in *S. cerevisiae* are associated with the production of biofuels (mainly ethanol and butanediol). One exception is the pair *ARO4*/*ARO7*, which catalyze the first committed step of the AAA pathway and the intermediate step, *chorismate mutase*. This design has been used in the production of specialized compounds such as resveratrol, styrene and 4-amino-benzoate. Another exception is the pair IDI1/ERG20 that catalyze different steps of terpenoid biosynthesis and is commonly up-regulated for the production of mono-terpenoids.

## Discussion

In this work, we used text-mining to create a database of experimentally validated strain design strategies. This extends the work performed over a decade ago by Wrinkle and co-workers ^24,25^. However, whereas their database contained hundreds of manually curated entries for two organisms, we opted to sacrifice curation for coverage and developed a fully automated pipeline. Our database contains over 15.000 research articles, covering 25 years of scientific literature, spanning a wide variety of microbial hosts and target compounds.

Our text mining workflow comprises several steps, including: automated queries on PubMed, data scraping techniques for extraction of metadata, machine learning for article classification, natural language processing (NLP) for detecting target compounds and modified genes ^49,50^. We used the industry-standard NLP tool, SpaCy ^42,51^, for several tasks such as parsing, tokenization, stemming and part-of-speech tagging. This automation will allow regularly updating the database with the most recent literature and generating new releases. One current limitation in our work is the inability to distinguish gene modifications as part of multiple strain design strategies implemented in the same publication and to compare their expected performance (some publications contain information on successful as well as unsuccessful strain design attempts). We were also unable to retrieve information on experimental conditions (such as growth media, temperature and pH) or production rates/yields/titers, as these are not systematically represented in a consistent format (i.e., they can be part of text, tables, figures, etc.). However, with the recent explosion in generative AI and large language models (LLMs), this may soon become feasible.

By analyzing data from thousands of successful case studies we can assess the effectiveness of various genetic modifications in different organisms, which can guide the design of new microbial cell factories ^52–54^. Looking at the association between microbial hosts and target products, we observe the utilization of generalist hosts (like *E. coli* and *S. cerevisiae*), which have been used to make all kinds of compounds, as well as specialist hosts (like *K. pastoris, Y. lipolytica*, and *C. glutamicum*), which are optimized for a given class of compounds. The distribution of hosts over the last few years reveals a recent trend in the preference for specialized hosts, most likely associated with the development of suitable genetic toolboxes for these microbes.

Despite the wide variety of hosts and target products across the thousands of publications, we observe a strong conservation at the level of genes that are frequently manipulated. Not surprisingly, these are genes encoding for enzymes in central carbon metabolism, mainly in the vicinity of the split between upper glycolysis and the pentose-phosphate pathway, and around the pyruvate node (TCA cycle, anaplerotic reactions, and fermentative pathways). Interestingly, these genes can just equally be targets for overexpression or deletion/under-expression. Therefore, it seems that during several decades of metabolic engineering, the main regulatory points for metabolic control have been identified. In other words, we know which buttons to press, the main challenge is knowing how to finely turn the knobs. The co-occurrence of targeted genes across different experimental designs reveals a high degree of modularity, i.e., depending on the desired product, there are small clusters of genes that are frequently manipulated together. This modularity is consistent with the concept, proposed by several authors, of building microbial chassis strains optimized to produce the precursors of different classes of compounds ^55–57^.

Overall, our database provides a stepping stone to start a new strain design project. It can guide the selection of a suitable host for the selected product and the exploration of experimentally validated genetic interventions. By building upon already proven and tested designs, metabolic engineers can try to improve their strains through further design-build-test-learn cycles, without reinventing the wheel at the start of every project.

## Acknowledgements

This work was funded by the Norwegian Research Council (SFI - Industrial Biotechnology, project 309558).

## Supplementary Figures

**Supplementary Figure 1.**
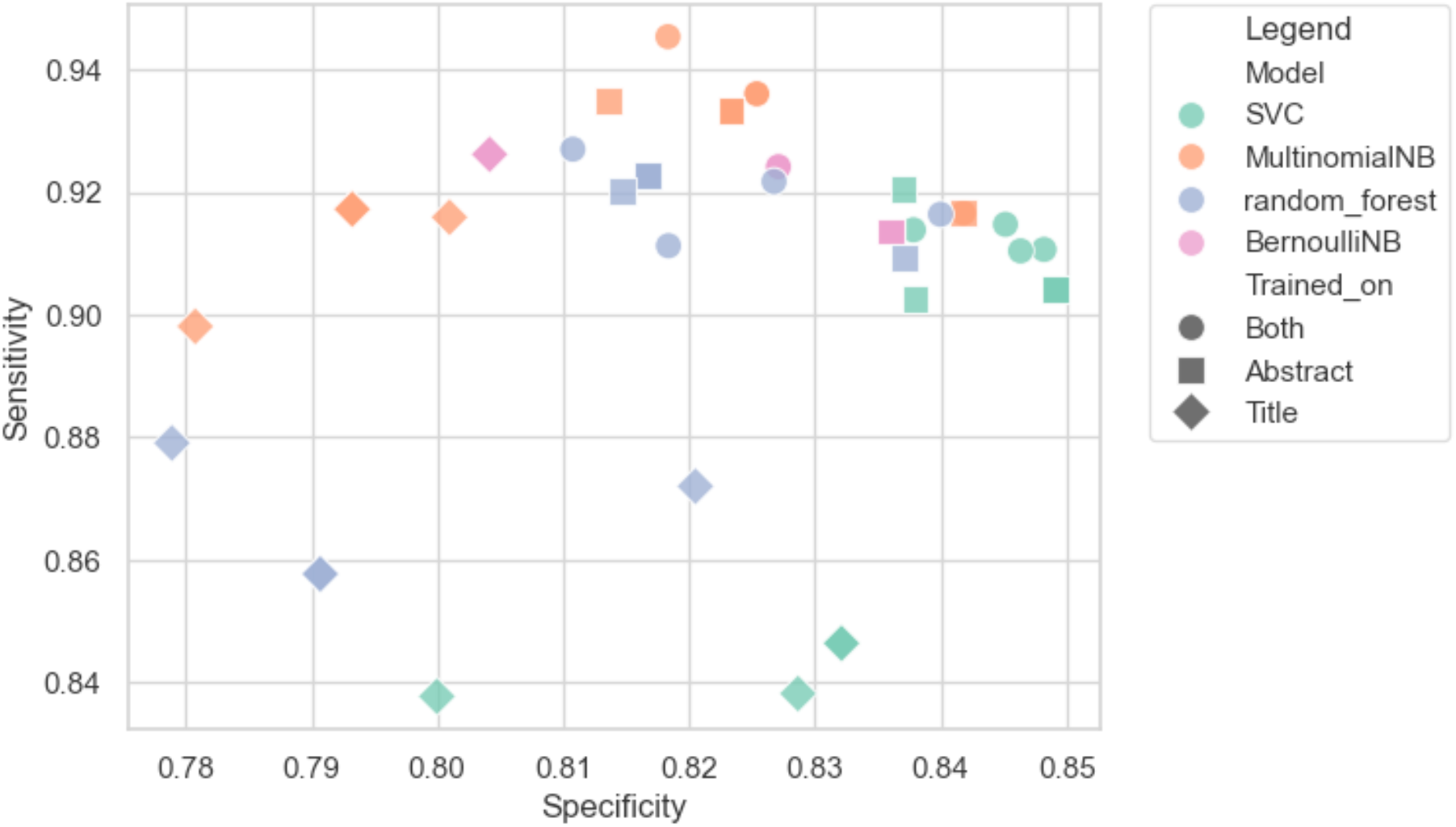
Specificity and sensitivity of the classification algorithms: Random Forests (RF), Bernoulli Naive Bayes (BNB), Support Vector Classifier (SVC), and Multinomial Naive Bayes (MNB).

## Notes

### Competing Interest Statement

The authors have declared no competing interest.

## References

1. Ko, Y.-S. et al. Tools and strategies of systems metabolic engineering for the development of microbial cell factories for chemical production. Chem. Soc. Rev. 49, 4615–4636 (2020).

2. Yang, D., Eun, H., Prabowo, C. P. S., Cho, S. & Lee, S. Y. Metabolic and cellular engineering for the production of natural products. Curr. Opin. Biotechnol. 77, 102760 (2022).

3. Gao, C. et al. Advances in microbial engineering for the production of value-added products in a biorefinery. Syst. Microbiol. Biomanufacturing 3, 246–261 (2023).

4. Gendoo, D. M. A. Overview of Bioinformatics Software and Databases for Metabolic Engineering. in Computational Biology and Machine Learning for Metabolic Engineering and Synthetic Biology (ed. Selvarajoo, K.) vol. 2553 265–274 (Springer US, New York, NY, 2023).

5. Adegboye, M. F., Ojuederie, O. B., Talia, P. M. & Babalola, O. O. Bioprospecting of microbial strains for biofuel production: metabolic engineering, applications, and challenges. Biotechnol. Biofuels 14, 5 (2021).

6. Hans, S. et al. Rebooting life: engineering non-natural nucleic acids, proteins and metabolites in microorganisms. Microb. Cell Factories 21, 100 (2022).

7. Kim, G. B., Choi, S. Y., Cho, I. J., Ahn, D.-H. & Lee, S. Y. Metabolic engineering for sustainability and health. Trends Biotechnol. 41, 425–451 (2023).

8. Nielsen, J. & Keasling, J. D. Engineering Cellular Metabolism. Cell 164, 1185–1197 (2016).

9. Carbonell, P. et al. An automated Design-Build-Test-Learn pipeline for enhanced microbial production of fine chemicals. Commun. Biol. 1, 66 (2018).

10. Tafur Rangel, A. E. et al. In silico Design for Systems-Based Metabolic Engineering for the Bioconversion of Valuable Compounds From Industrial By-Products. Front. Genet. 12, 633073 (2021).

11. Carbonell, P. Synthetic biology design tools for metabolic engineering. in Microbial Cell Factories Engineering for Production of Biomolecules 65–77 (Elsevier, 2021). doi:10.1016/B978-0-12-821477-0.00005-2.

12. Sveshnikova, A., MohammadiPeyhani, H. & Hatzimanikatis, V. Computational tools and resources for designing new pathways to small molecules. Curr. Opin. Biotechnol. 76, 102722 (2022).

13. Delépine, B., Duigou, T., Carbonell, P. & Faulon, J.-L. RetroPath2.0: A retrosynthesis workflow for metabolic engineers. Metab. Eng. 45, 158–170 (2018).

14. Sulheim, S., Fossheim, F. A., Wentzel, A. & Almaas, E. Automatic reconstruction of metabolic pathways from identified biosynthetic gene clusters. BMC Bioinformatics 22, 81 (2021).

15. Otero-Muras, I. & Carbonell, P. Automated engineering of synthetic metabolic pathways for efficient biomanufacturing. Metab. Eng. 63, 61–80 (2021).

16. Hartmann, A. et al. OptPipe - a pipeline for optimizing metabolic engineering targets. BMC Syst. Biol. 11, 143 (2017).

17. Shen, F. et al. OptRAM: In-silico strain design via integrative regulatory-metabolic network modeling. PLOS Comput. Biol. 15, 1006835 (2019).

18. Han, T., Nazarbekov, A., Zou, X. & Lee, S. Y. Recent advances in systems metabolic engineering. Curr. Opin. Biotechnol. 84, 103004 (2023).

19. Kanehisa, M. KEGG: Kyoto Encyclopedia of Genes and Genomes. Nucleic Acids Res. 28, 27–30 (2000).

20. The UniProt Consortium et al. UniProt: the Universal Protein Knowledgebase in 2023. Nucleic Acids Res. 51, D523–D531 (2023).

21. Caspi, R. et al. The MetaCyc database of metabolic pathways and enzymes - a 2019 update. Nucleic Acids Res. 48, D445–D453 (2020).

22. Chang, A. et al. BRENDA, the ELIXIR core data resource in 2021: new developments and updates. Nucleic Acids Res. 49, D498–D508 (2021).

23. Nakazawa, S. et al. History-Driven Genetic Modification Design Technique Using a Domain-Specific Lexical Model for the Acceleration of DBTL Cycles for Microbial Cell Factories. ACS Synth. Biol. 10, 2308–2317 (2021).

24. Winkler, J. D., Halweg-Edwards, A. L. & Gill, R. T. The LASER database: Formalizing design rules for metabolic engineering. Metab. Eng. Commun. 2, 30–38 (2015).

25. Winkler, J. D., Halweg-Edwards, A. L. & Gill, R. T. Quantifying complexity in metabolic engineering using the LASER database. Metab. Eng. Commun. 3, 227–233 (2016).

26. Database resources of the National Center for Biotechnology Information. Nucleic Acids Res. 42, D7–D17 (2014).

27. The Europe PMC Consortium. Europe PMC: a full-text literature database for the life sciences and platform for innovation. Nucleic Acids Res. 43, D1042–D1048 (2015).

28. Achakulvisut, T., Acuna, D. & Kording, K. Pubmed Parser: A Python Parser for PubMed Open-Access XML Subset and MEDLINE XML Dataset XML Dataset. J. Open Source Softw. 5, 1979 (2020).

29. Jenuth, J. P. The NCBI: Publicly Available Tools and Resources on the Web. in Bioinformatics Methods and Protocols vol. 132 301–312 (Humana Press, New Jersey, 1999).

30. Ostell, J. M. Entrez: The NCBI Search and Discovery Engine. in Data Integration in the Life Sciences (eds. Bodenreider, O. & Rance, B.) vol. 7348 1–4 (Springer Berlin Heidelberg, Berlin, Heidelberg, 2012).

31. Sayers, E. W. et al. Database resources of the national center for biotechnology information. Nucleic Acids Res. 50, D20–D26 (2022).

32. Degtyarenko, K. et al. ChEBI: a database and ontology for chemical entities of biological interest. Nucleic Acids Res. 36, D344–D350 (2007).

33. Hastings, J. et al. ChEBI in 2016: Improved services and an expanding collection of metabolites. Nucleic Acids Res. 44, D1214–D1219 (2016).

34. Kanehisa, M. Toward understanding the origin and evolution of cellular organisms. Protein Sci. 28, 1947–1951 (2019).

35. Kanehisa, M., Furumichi, M., Sato, Y., Kawashima, M. & Ishiguro-Watanabe, M. KEGG for taxonomy-based analysis of pathways and genomes. Nucleic Acids Res. 51, D587–D592 (2023).

36. Fraile Navarro, D. et al. Clinical named entity recognition and relation extraction using natural language processing of medical free text: A systematic review. Int. J. Med. Inf. 177, 105122 (2023).

37. Cock, P. J. A. et al. Biopython: freely available Python tools for computational molecular biology and bioinformatics. Bioinformatics 25, 1422–1423 (2009).

38. Pedregosa, F. et al. Scikit-learn: Machine Learning in Python. J. Mach. Learn. Res. 12, 2825–2830 (2011).

39. Salton, G. & Buckley, C. Term-weighting approaches in automatic text retrieval. Inf. Process. Manag. 24, 513–523 (1988).

40. Dhakal, K. Unpaywall. J. Med. Libr. Assoc. 107, (2019).

41. pdfminer.six contributors & pyHanko. pdfminer.six. GitHub (2025).

42. Honnibal, M., Montani, I., Van Landeghem, S. & Boyd, A. spaCy: Industrial-strength Natural Language Processing in Python. https://doi.org/10.5281/zenodo.1212303 (2020) doi:10.5281/zenodo.1212303.

43. Moretti, S. et al. MetaNetX/MNXref – reconciliation of metabolites and biochemical reactions to bring together genome-scale metabolic networks. Nucleic Acids Res. 44, D523–D526 (2016).

44. King, Z. A. et al. BiGG Models: A platform for integrating, standardizing and sharing genome-scale models. Nucleic Acids Res. 44, D515–D522 (2016).

45. Deng, X., Li, Y., Weng, J. & Zhang, J. Feature selection for text classification: A review. Multimed. Tools Appl. 78, 3797–3816 (2019).

46. Xu, S. Bayesian Naïve Bayes classifiers to text classification. J. Inf. Sci. 44, 48–59 (2018).

47. Gan, S., Shao, S., Chen, L., Yu, L. & Jiang, L. Adapting Hidden Naive Bayes for Text Classification. Mathematics 9, 2378 (2021).

48. Kumar, A., Mitchener, J., King, Z. A. & Metallo, C. M. Escher-Trace: a web application for pathway-based visualization of stable isotope tracing data. BMC Bioinformatics 21, 297 (2020).

49. Lewinski, N. A. & McInnes, B. T. Using natural language processing techniques to inform research on nanotechnology. Beilstein J. Nanotechnol. 6, 1439–1449 (2015).

50. Oikonomou, E. D. et al. How natural language processing derived techniques are used on biological data: a systematic review. Netw. Model. Anal. Health Inform. Bioinforma. 13, 23 (2024).

51. Amade, D., Chandra, R., Sinha, V. K. & Anand, D. Automatic Text Summarization Using NLTK & Spacy*. SSRN Electron. J. https://doi.org/10.2139/ssrn.4742012 (2024) xdoi:10.2139/ssrn.4742012.

52. Carbonell, P., Radivojevic, T. & García Martín, H. Opportunities at the Intersection of Synthetic Biology, Machine Learning, and Automation. ACS Synth. Biol. 8, 1474–1477 (2019).

53. Presnell, K. V. & Alper, H. S. Systems Metabolic Engineering Meets Machine Learning: A New Era for Data-Driven Metabolic Engineering. Biotechnol. J. 14, 1800416 (2019).

54. Roy, S. et al. Multiomics Data Collection, Visualization, and Utilization for Guiding Metabolic Engineering. Front. Bioeng. Biotechnol. 9, 612893 (2021).

55. Matsumoto, T., Tanaka, T. & Kondo, A. Engineering metabolic pathways in Escherichia coli for constructing a “microbial chassis” for biochemical production. Bioresour. Technol. 245, 1362–1368 (2017).

56. Jouhten, P. et al. Yeast metabolic chassis designs for diverse biotechnological products. Sci. Rep. 6, 29694 (2016).

57. Liu, J., Wu, X., Yao, M., Xiao, W. & Zha, J. Chassis engineering for microbial production of chemicals: from natural microbes to synthetic organisms. Curr. Opin. Biotechnol. 66, 105–112 (2020).

